# Statistical BURST imaging for high-fidelity biomolecular ultrasound

**DOI:** 10.64898/2026.03.13.711726

**Authors:** Sunho Lee, Shirin Shivaei, Mikhail G. Shapiro

## Abstract

Ultrasound is emerging as a method for molecular and cellular imaging by connecting the versatile physics of sound waves to protein-based contrast agents such as gas vesicles (GVs). BURST is a common imaging mode that leverages the strong, transient echoes generated when GVs collapse under acoustic pressure to enable highly sensitive ultrasound visualization of cells and biomolecules, down to the single cell level. However, BURST is vulnerable to fluctuating background signals, with large-amplitude fluctuations in scattering, as often present *in vivo*, obscuring genuine GV responses. In this study, we mathematically examine this limitation and show that incorporating statistical metrics such as correlation or temporal contrast-to-noise ratio effectively suppresses unwanted non-GV voxels and quantifies detection confidence, including in image sequences in which GV collapse spans multiple frames. Compared with prior methods, our approach enhances the clarity of BURST images and provides probabilistic interpretations of GV signals, facilitating more reliable analysis of ambiguous *in vivo* molecular imaging, as we demonstrate in imaging tumor-homing probiotics and gene expression in the brain.

---

Ultrasound imaging, long established as a modality for visualizing anatomy and physiology, is emerging as a method for deep tissue molecular and cellular imaging thanks to the development of genetically encodable contrast agents. In particular, gas vesicles (GVs), air-filled protein nanostructures derived from certain species of bacteria and archaea^1^, have emerged as versatile agents that link cellular and biomolecular activity to ultrasound signal^2^. After the discovery that GVs produce nonlinear acoustic responses to ultrasound^3^, two non-linear imaging strategies have become common: amplitude modulation (AM)^4,5^ and BURST^6^, the latter offering sensitivity down to the single-cell level. In BURST, the transmitted ultrasound pressure is raised above the GV collapse threshold during image acquisition, causing the GVs to irreversibly collapse and generate strong transient signals that are subsequently separated from static background by post-processing of the image sequence (**Fig. 1a-b**). Although it requires collapsing the GVs, the sensitivity of BURST has been pivotal to enable several *in vivo* applications of biomolecular ultrasound imaging^7–9^.

**FIG. 1.**
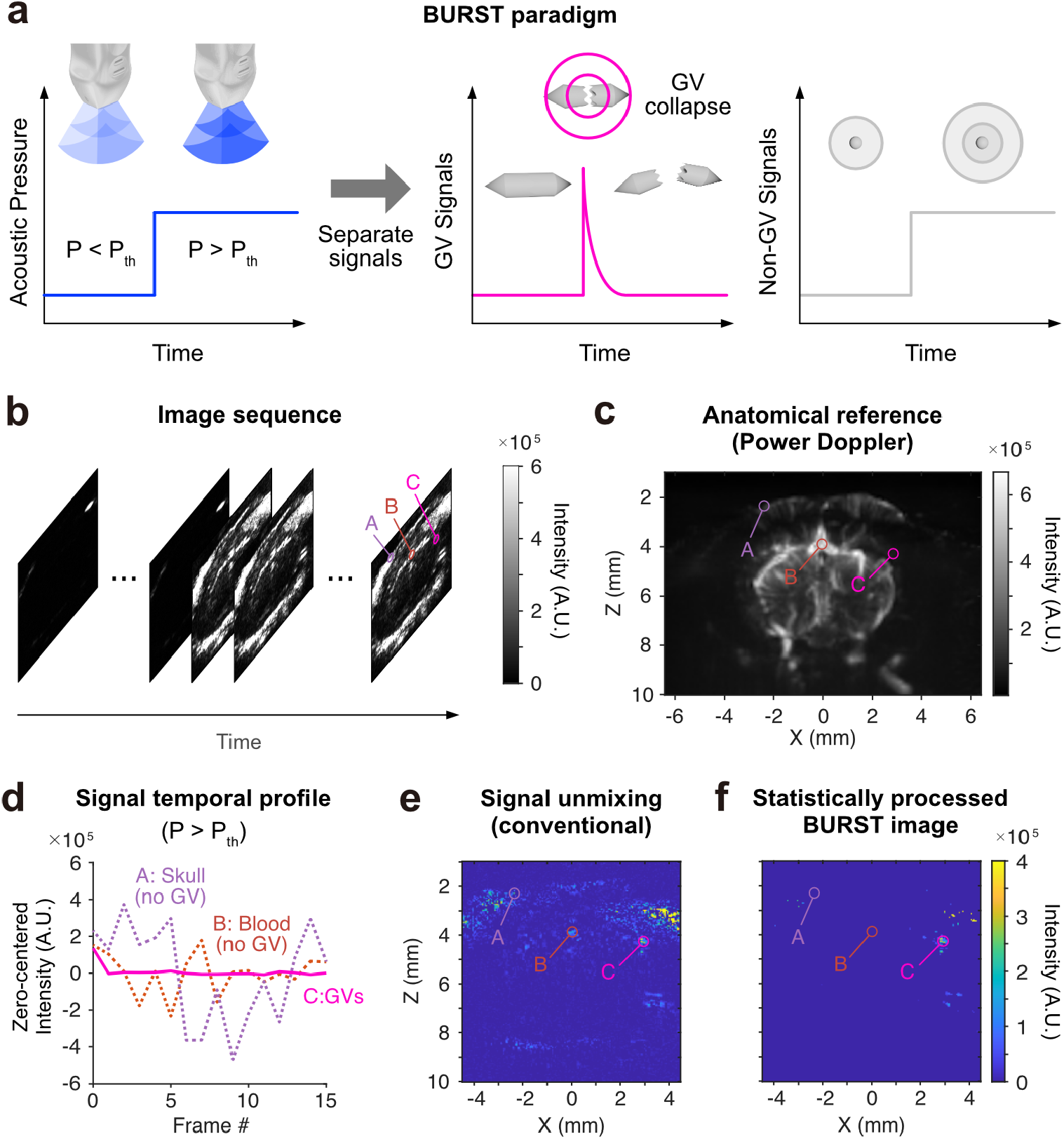
Susceptibility of conventional BURST processing to background fluctuations. (a) BURST paradigm. When ultrasound is transmitted at acoustic pressures above the collapse threshold, GVs generate strong, transient signals, whereas non-GV structures backscatter at a constant level. (b) Example BURST image sequence from *Shivaei et al*.8 (AM-mode). Voxels A/B/C: skull, blood vessels, GVs. (c) Power Doppler image acquired after GV collapse, shown for anatomical reference. (d) Temporal signal profiles at voxels A/B/C during high-pressure transmission. Non-GV signals are zero-centered using all 16 frames, while GV signals are processed using only the last 15 frames. (e) Final BURST image obtained using the conventional signal unmixing approach. (f) Final BURST image obtained using statistical processing.

Currently, two approaches are typically used to process BURST image sequences. The most intuitive is frame subtraction, in which a background frame is subtracted from the frame containing transient GV collapse signal^7,9,10^. A more sophisticated technique, first formalized in the original BURST imaging paper^6^, is template-based signal unmixing, which disassembles each voxel’s time course into GV, background, and offset components by fitting predefined temporal templates in a matrix formulation equivalent to a general linear model (GLM).

However, both frame subtraction and template-based unmixing are vulnerable to large background fluctuations, which cause non-GV voxels in BURST images to exhibit intensities proportional to the fluctuation level (**Fig. 1c-e**). This is especially problematic when imaging low-density or sparsely distributed GVs *in vivo*, where reflections or clutter from tissue may appear as prominently as the GV signal in the final BURST image. Moreover, since the effect of tissue nonlinearity increases with acoustic pressure – and BURST employs high pressures (2-4 MPa^11^) – these tissue backgrounds are not always adequately suppressed, even with AM-mode BURST. In this paper, we show that incorporating statistical information into the signal unmixing paradigm, analogously to statistical parametric mapping (SPM) in functional magnetic resonance imaging (fMRI) GLM analyses, effectively addresses the background fluctuation problem (**Fig. 1f, Fig. 2**). First, we relate template-based unmixing to frame subtraction and explain why subtraction fails under large fluctuations. We then demonstrate that incorporating statistics of the background and GV signals via correlations or temporal contrast-to-noise ratio (tCNR) substantially improves BURST images and provides useful probabilistic interpretations of the signals. Lastly, we extend the method to cases in which GV collapse spans multiple frames. This paper also justifies our use of this method in *Shivaei et al*.^8^. For simplicity, we assume 1) negligible motion on the timescale of image sequence acquisition, 2) spatial/temporal independence across voxels, and 3) Gaussian fluctuations. These assumptions are further discussed at the end of this paper.

**FIG. 2.**
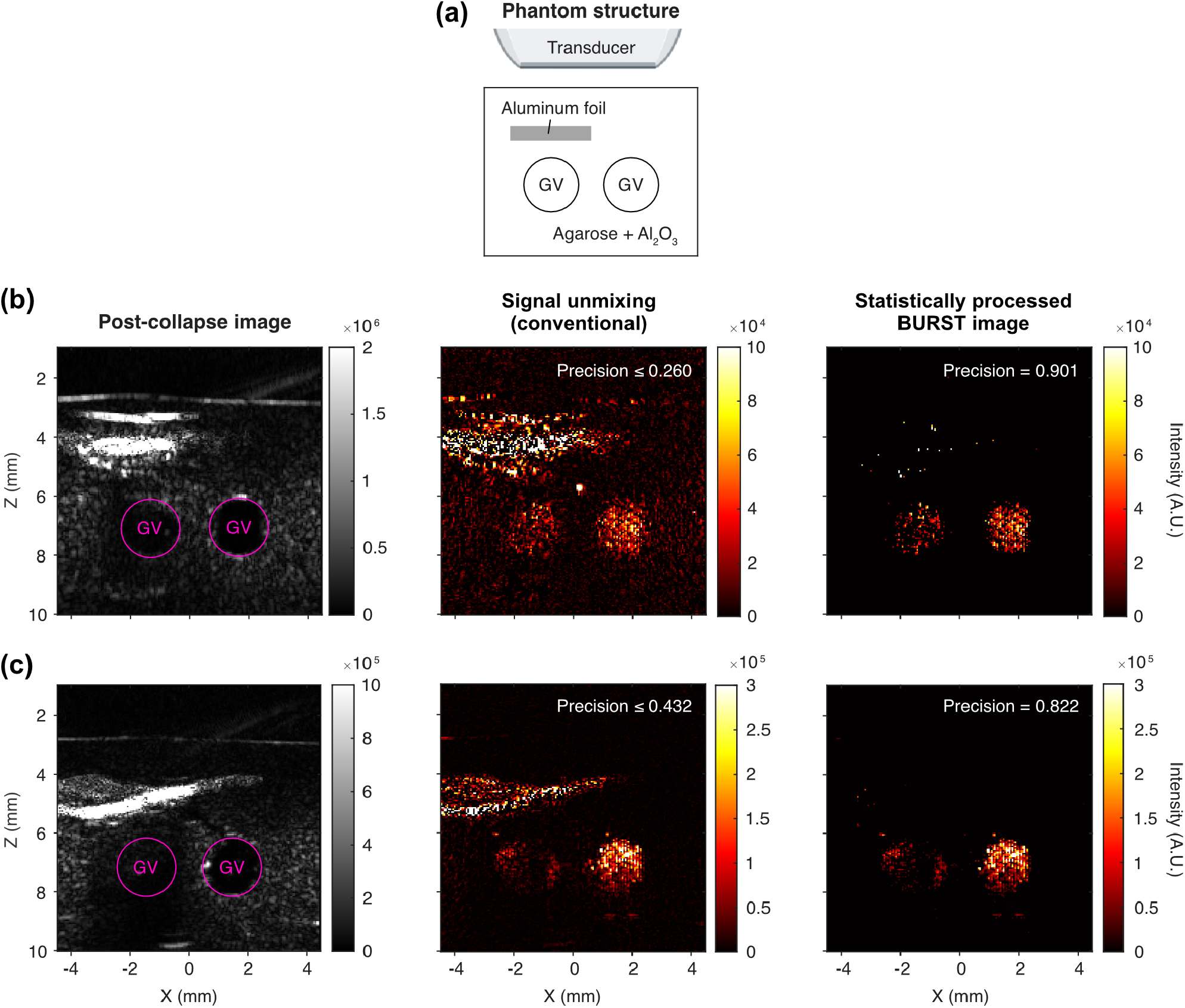
Statistical processing of *in vitro* BURST images. (a) Structure of tissue-mimicking GV phantoms. The background consists of 1% agarose + 0.2% Al2O3, and GVs (optical density = 0.4) are embedded in two wells of 0.5% agarose. To generate strong artifacts, aluminum foil is embedded above the GV wells. Foil thickness is exaggerated in the schematic. (b-c) Example BURST images. The last post-collapse frames are shown to visualize phantom structure. For statistical processing, *p* = 1× 10-^4^ is used. The number of post-collapse frames is 99 for (b) and 19 for (c). Precision is defined as the fraction of classified-positive pixels that fall within the GV wells (i.e., TP / (TP + FP)). For BURST images with signal unmixing, the optimal precision values are shown.

To better understand why the signal unmixing method is vulnerable to strong background fluctuations, we first rewrite its matrix equation in a subtraction form. Let **X** be the design matrix whose rows are the template vectors for GV collapse, background, and offset signals:

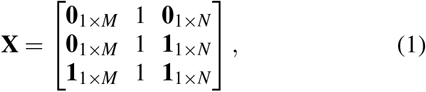

where *M* is the number of pre-collapse frames, *N* is the number of post-collapse frames (no GVs exist), and collapse occurs at frame *M* + 1. The matrices **0**_*a*_×_*b*_ and **1**_*a*_ ×_*b*_ denote the zero and all-ones matrices of size *a* × *b*, respectively. Given a sequence of ultrasound images **Y** = **F**_pre,1_ · · · **F**_pre,*M*_ **F**_collapse_ **F**_post,1_ · · · **F**_post,*N*_ ∈ ℝ^*k*×(*M*+*N*+1)^ where each **F**_(_ ·, · _)_ ∈ ℝ^*k*^ is a flattened image over *k* voxels, the matrix equation is:

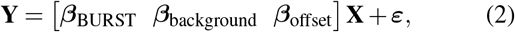

where the parameters ***β***_(·)_∈ ℝ^*k*^ and voxel-dependent fluctuations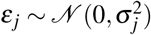, where *N* denotes a normal distribution and 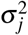is the variance for voxel *j*. Using the least-squares solution (**Appendix A 1**), the estimator of the GV component simplifies to

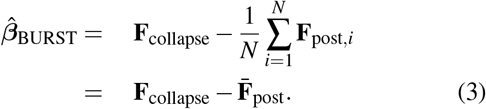

Therefore, template-based unmixing is basically equivalent to a subtraction method: it estimates GV signals by subtracting the mean of the post-collapse frames (i.e., the estimated background at high pressure) from the collapse frame. Note that our only interest,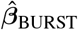, does not depend on any pre-collapse frames. The design matrix (**Eq. 1**) can thus be reduced to

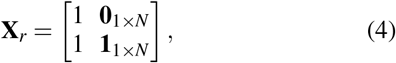

and **Eq. 2** becomes

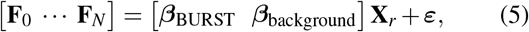

where for notational convenience we define **F**_0_ = **F**_collapse_ and **F**_*i*_ = **F**_post,*i*_ (*i >* 0). Hence, for each voxel, the multiple linear regression reduces to a simple linear regression.

At a background voxel *j* with no GVs, the intensities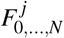are independently and identically distributed as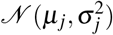, where *µ*_*j*_ denotes the mean intensity. Then, from **Eq. 3**,

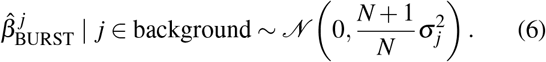

**Eq. 6** implies that subtraction-based methods, including template-based unmixing, can yield large intensities in the BURST image at purely background voxels when temporal fluctuations are strong – even in the absence of GVs. This issue is especially pronounced near tissue interfaces, where reflections and clutter artifacts dominate. Since BURST employs high pressure and the influence of tissue nonlinearity increases with pressure, such backgrounds can persist even with AM mode and may remain confounding, especially when GV expression is weak (**Fig. 1b-e**).

BURST images derived from template-based unmixing 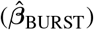do not explicitly reflect the degree of similarity between each voxel’s time course and the GV signal template **T** = [1 **0**_1_× _*N*_], but this can be readily measured through statistical methods. Specifically, the statistical significance of simple linear regression (**Eq. 5**) can be calculated via the correlation coefficient *r*_*BURST*_:

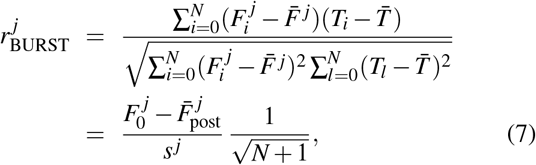

where 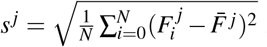is the sample standard deviation of voxel *j*’s signal over the collapse and post-collapse frames. Because **T** = [1 **0**_1_×_*N*_], the correlation reduces to the subtraction form 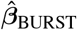(**Eq. 3**) divided by the voxel’s sample standard deviation *s* ^*j*^ with an appropriate scaling factor. This standardization suppresses voxels with strong fluctuations that otherwise appear as high-intensity artifacts in subtraction-based BURST images (**Fig. 3a-e**). For background voxels, the sampling distribution of *r*_*BURST*_ is^12,13^

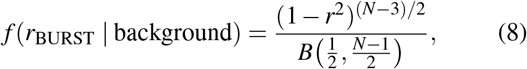

where *B*(·, ·) is the beta function. Notably, this distribution depends only on the number of frames *N* + 1, not on voxel-specific parameters (e.g., *σ*_*j*_), implying that the correlation values of background voxels across the entire image fall along the same distribution (**Fig. 3a,d**; see **Supplementary Material** for further discussion of the distribution of *r*_*BURST*_ in the real image dataset). Since **Eq. 8** is cumbersome, correlation is typically transformed into a *t*-statistic with *N* − 1 degrees of freedom^12,13^:

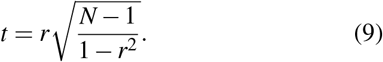

**FIG. 3.**
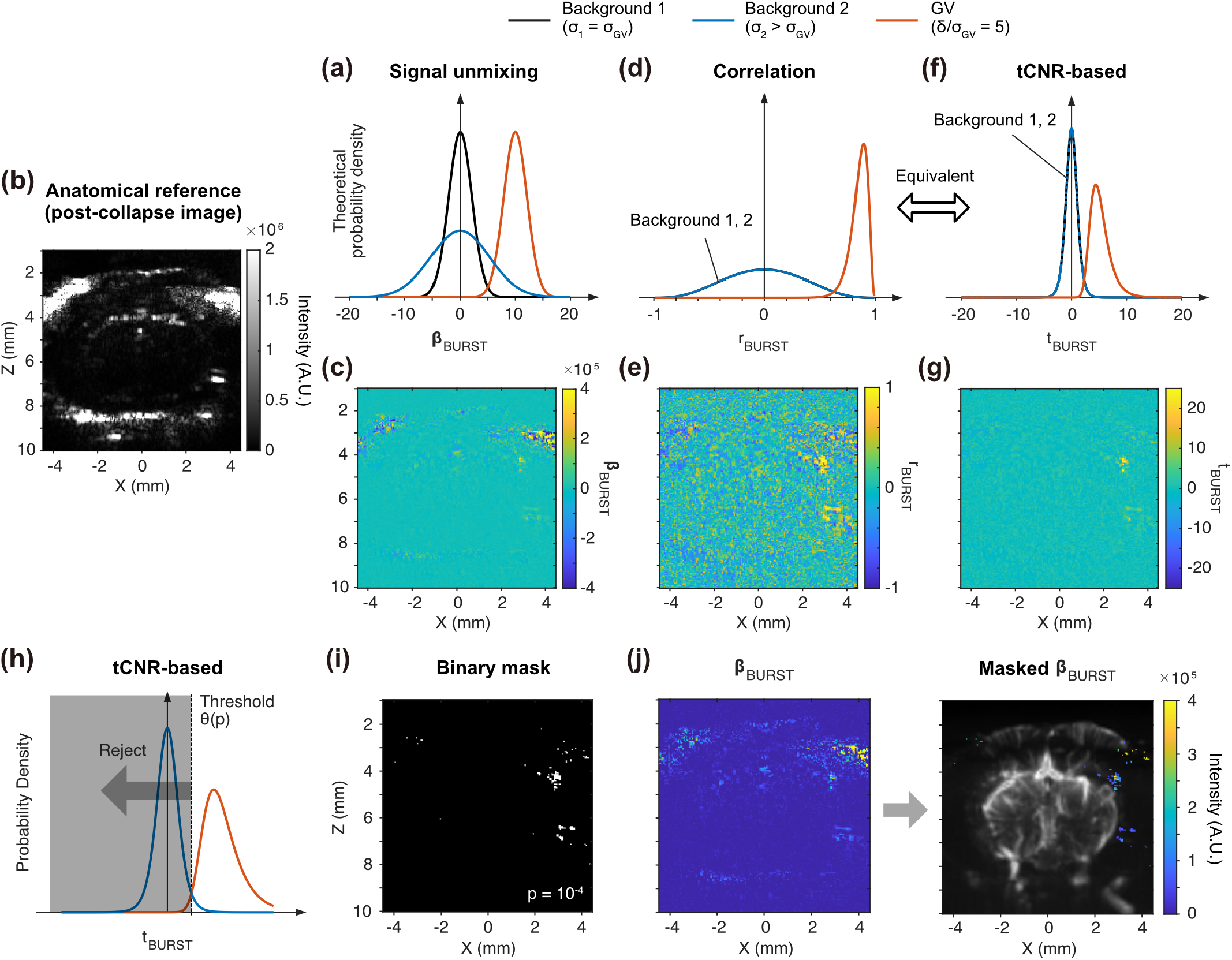
Statistical parameter maps of BURST and integration with intensity information. Theoretical probability density functions of (a) 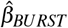, (d) *r*_*BURST*_, and (f) *t*_*BURST*_ at background and GV pixels. Background 1 has the same level of fluctuation (*σ*) as the GV pixel, whereas Background 2 has larger fluctuations. Parameters used in these plots: *σ*_1_ = *σGV* = 2, *σ*_2_ = 5, *δ* = 10, and *N* = 9. (b) Post-collapse image for anatomical reference and example maps of (c) 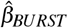(e) *r*_*BURST*_, and (g) *t*_*BURST*_ are shown using data from *Shivaei et al*.^8^. Negative 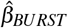values are retained (rather than zeroed) to allow better comparison across maps, although conventional signal unmixing typically sets them to zero. (h) A *t*_*BURST*_ map can be thresholded at a desired p-value to suppress insignificant background voxels. (i) Example binary mask from thresholding *t*_*BURST*_. *p* = 1 × 10^−4^. (j) Applying the mask to the intensity map *β*_*BURST*_ merges the statistical and intensity information to produce the final image. The final BURST image is overlaid with a power Doppler image.

Meanwhile, observing the fraction 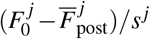in **Eq. 7**, an alternative statistical measure can be considered. Suppose we standardize the subtraction in the numerator using the sample standard d<u>eviation calculated only</u> from post-collapse frames, 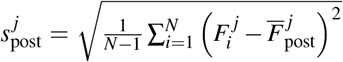, which more directly estimates background fluctuations than *s* ^*j*^. This metric, denoted as tCNR, with appropriate scaling,

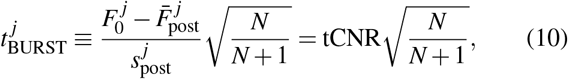

follows a *t*-distribution with *N−*1 degrees of freedom for background voxels.

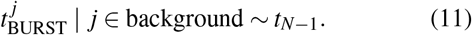

As demonstrated in **Appendix B**, this *t* ^*j*^ (**Fig. 3f-g**) is algebraically identical to the *t*-transformed correlation in **Eq. 9**.

Thus, *r*_BURST_ and *t*_BURST_ (i.e., tCNR-based) provide equivalent statistical representations of the BURST image, although *t*_BURST_ may better visualize significance due to the bounded range of *r*_BURST_ (**Fig. 3e,g**).

As in SPM for fMRI, one can threshold *t*_BURST_ or *r*_BURST_ maps using a desired *p*-value (false positive rate) to identify significant voxels likely to contain GVs (**Fig. 3h**). However, the raw BURST intensity itself 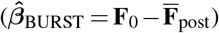also carries valuable information, such as reflectivity and backscattering strength. In fact, for background voxels with significantly small fluctuations—so small that their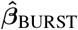 values are already well separated from those of GV voxels (consider a background distribution narrower than Background 1 in **Fig. 3a**)—standardization may rather reduce discriminability. To combine the strengths of both views, we create a binary mask from the thresholded *t*_BURST_ map (**Fig. 3i**) and apply it to the intensity map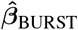 (**Fig. 3j**):

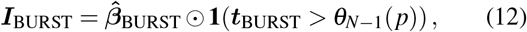

where ⊙ denotes element-wise multiplication, **1**(·) is the indicator function, and *θ*_*N*_*-*_1_(*p*) is the threshold from the cumulative distribution function (CDF) of the *t*-distribution with *N−* 1 degrees of freedom, corresponding to the chosen acceptance rate *p* (or equivalently, a *p*-value). By adjusting *p*, we can systemically suppress background voxels in the BURST intensity map 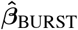(**Fig. 4a**).

**FIG. 4.**
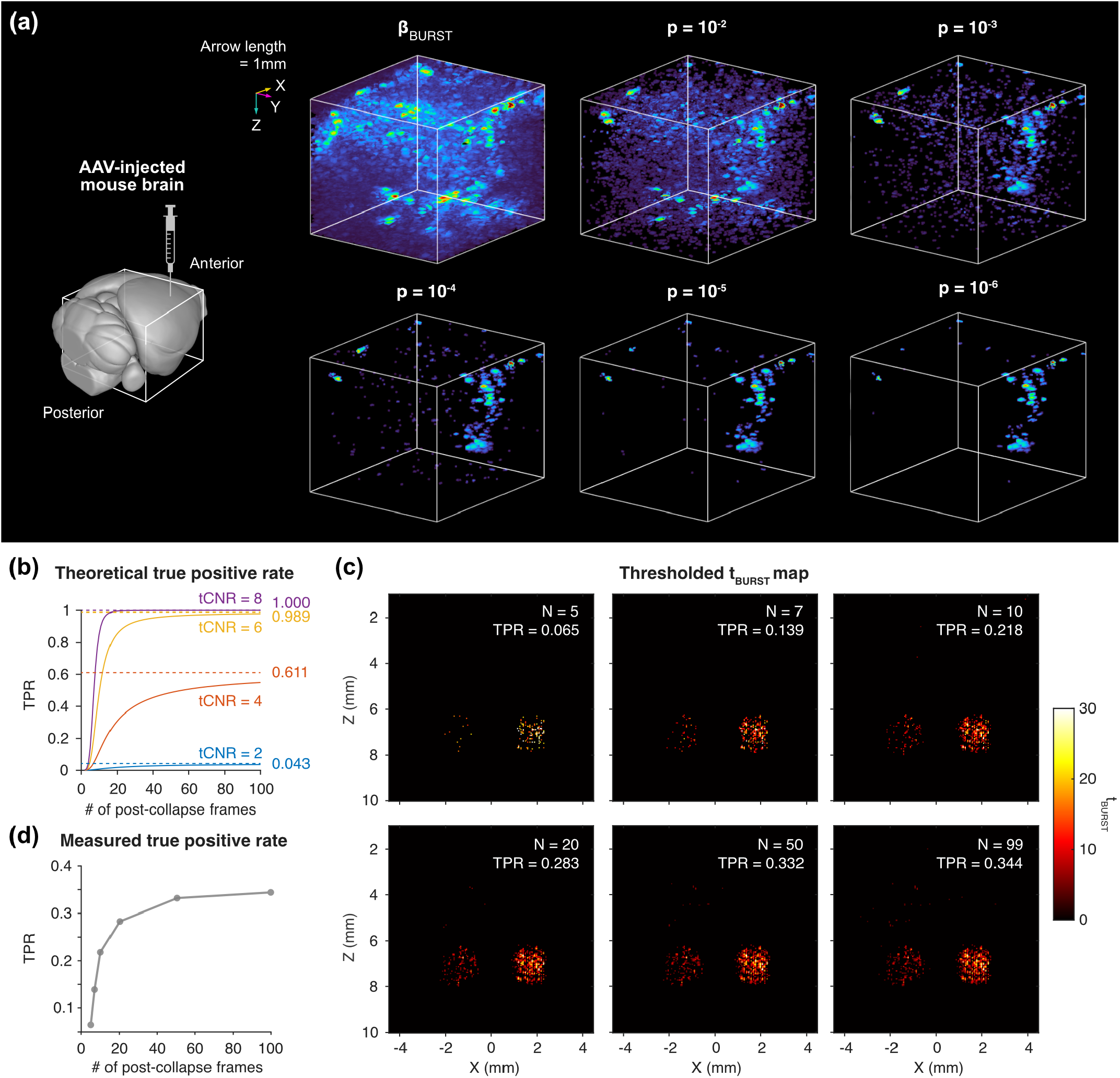
Effects of p-value selection and the number of post-collapse frames on BURST image processing. (a) Masked 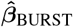maps are shown for various *p*-values using 3D B-mode-based BURST brain image data from *Shivaei et al*.^8^. The raw 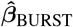 map is the final BURST image for the conventional signal unmixing method. Image dimensions are (*X,Y, Z*) = 8.4 ×7.5× 6.7 mm (171× 153 ×136 voxels). Colored arrows indicate 1 mm along each axis. 3D volumes are visualized with Napari^14^. (b) For each specified true tCNR (*δ*_GV_*/σ*) and number of post-collapse frames (*N*), the corresponding theoretical TPR values are plotted. Dashed lines indicate theoretical upper bounds on TPR. TPR is calculated when *p* = 1×10^−4^. (c) Thresholded *t*_BURST_ maps (*p* = 1 ×10^−4^) computed with varying numbers of post-collapse frames. Data are identical to **Fig. 2b**. TPR is measured as the fraction of positively classified pixels within GV wells (**Fig. 2a**). (d) Measured TPR as a function of the number of post-collapse frames.

While the ground-truth intensity of GV collapse is generally unknown, **Eq. 10** gives insight into the distribution of the resulting *t*_BURST_ values. Suppose for a GV voxel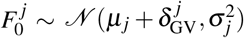and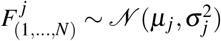,where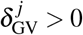is the collapse-induced impulse. Then its *t*_BURST_ follows a non-central *t*-distribution with *N* 1 degrees of freedom:

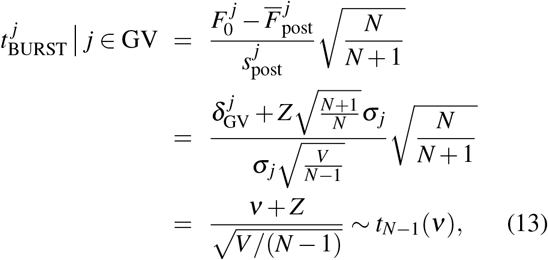

where *Z ∼* *N* (0, 1), 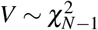, and the non-centrality param-eter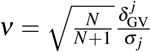. An estimate of the true tCNR 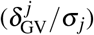 could be directly obtained as

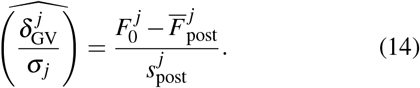

Since we now have the conditional distributions of *t*_BURST_ for both background and GV voxels, we can evaluate both the false positive and false negative rates for any voxel at a given threshold.

Given a GV’s tCNR, this framework also allows us to quantify how increasing the number of frames improves the classification performance and to derive its upper bound. In our context, a practical performance metric is the true positive rate (TPR), because we fix the true negative rate (TNR = 1 − *p*) with a desired *p*-value (**Fig. 3h**). As the degrees of freedom approach infinity (i.e., with infinitely many frames), the central and non-central *t*-distributions converge to *N* (0, 1) and*N* (*ν*, 1), respectively. The upper bound of the TPR can then be derived as

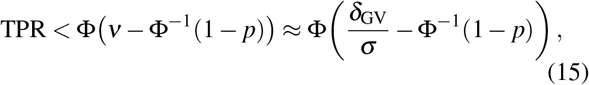

where Φ is the CDF of the standard normal distribution. The theoretical trends of TPR (**Fig. 4b**) and *in vitro* examples (**Fig. 4c-d**) illustrate that, as more post-collapse frames are used for processing BURST images, additional lower-tCNR voxels tend to appear after thresholding the statistical maps. These relationships help determine how many frames to collect for target BURST image quality, or, conversely, how much GV signal can be recovered under frame-limited conditions (e.g., due to tissue motion).

So far, we have focused on the case of a single collapse frame. In practice, however, GV collapse can span multiple frames – often exhibiting exponentially decaying, rather than spike-like, signals – or occur later than the first high-pressure frame, depending on ultrasound parameters (e.g., high GV concentration, low acoustic pressure causing shielding effects) and experimental context (e.g., tissue attenuation)^6,8,9^.

Assuming we can determine after which frame no further GV collapse occurs (e.g., by inspecting the image sequence), we can treat each collapse frame as an independent collapse event and construct a design matrix as following:

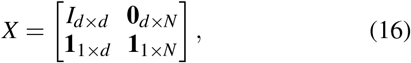

where *d* is the number of collapse frames. The BURST in tensity image corresponding to each collapse frame is then estimated as (**Appendix A 2**)

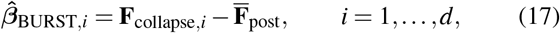

which mirrors the form of **Eq. 3** applied to each collapse frame individually. For each 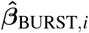 a statistical map *t*_BURST,*i*_ can be computed and combined with **Eq. 12** (**Fig. 5a-b**). The final BURST image is obtained by summing all masked components (**Fig. 5c**):

**FIG. 5.**
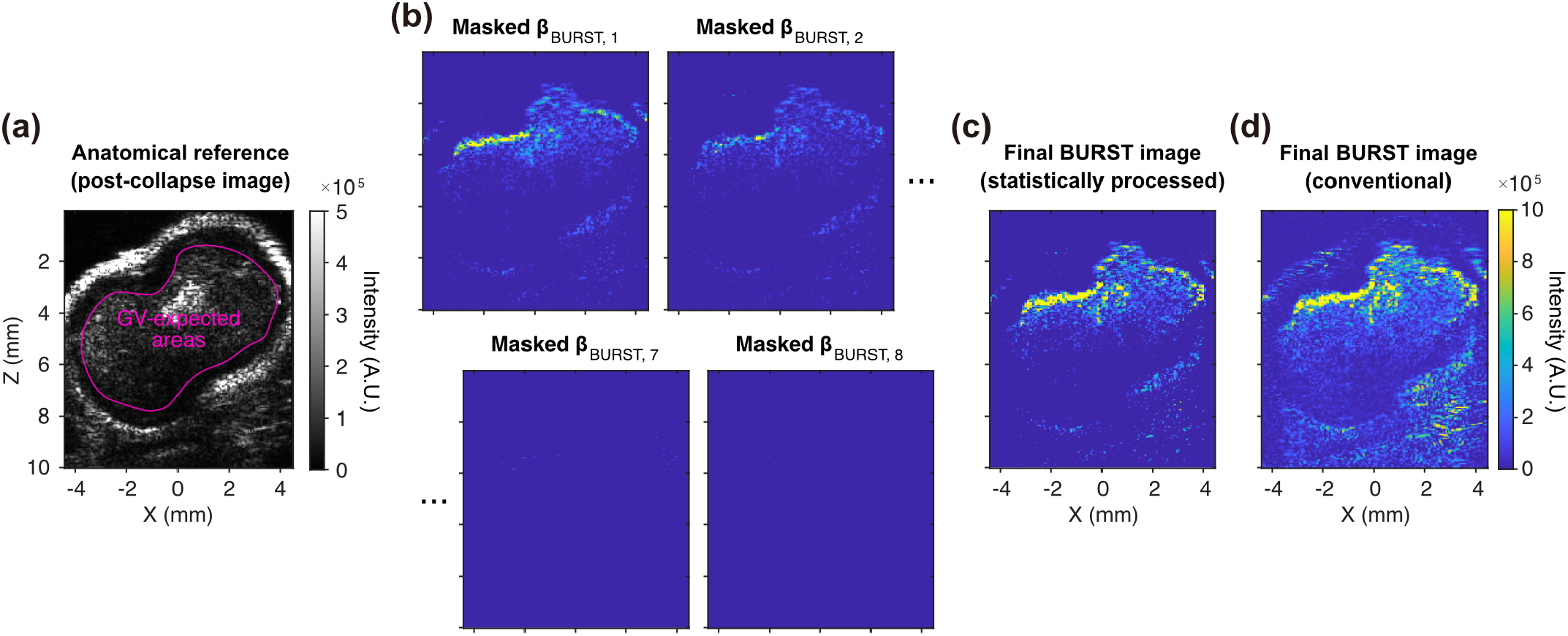
BURST processing when GV collapse spans multiple frames. (a) Post-collapse image of subcutaneous tumor shown for anatomical reference. Data from *Hurt et al*.^11^. (b) Example masked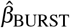 maps for each collapse frame (*d* = 8) in AM-based BURST imaging data. *p* = 1 ×10^−4^. (c) Final statistical BURST image obtained by summing all statistically masked components. (d) Final BURST image obtained by summing 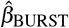 maps without statistical masks. For all summations, negative values in each 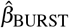 map are zeroed.

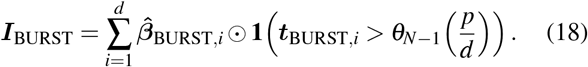

Note that the Bonferroni correction is applied to the *p*-value to compensate for the multiple comparisons introduced by the *d* collapse frames. Compared to summing the 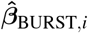 maps with zeroing negative values as done in conventional methods (**Fig. 5d**), the statistical method suppresses high-intensity voxels in background tissue more effectively.

In summary, we showed that subtraction-based BURST processing, including the originally proposed template-based signal unmixing^6^ (a GLM formulation), is inevitably vulnerable to regions with strong temporal fluctuations, which inflate background-voxel intensities and obscure genuine GV responses in the final BURST images. By incorporating statistical information, in a manner similar to SPM in fMRI analysis, we were able to suppress non-GV regions and improve both the quality and interpretability of BURST images. Using correlation or tCNR-based maps, which are statistically equivalent, we could set thresholds to filter out non-significant voxels and estimate type I/II errors for each voxel based on their conditional distributions. This framework was also further extended to cases where GV collapse occurs across multiple frames. We demonatrated the advantages of this approach in images acquired in imaging GV expression in the brain and visualizing tumor-homing probiotics in mice.

Although the current approach is simple and likely the most efficient for practical applications, more precise statistical inference is possible under complicated but more realistic assumptions. For instance, we grounded our derivations in Gaussian temporal fluctuations, yet the envelope of ultrasound signals is often better modeled by non-Gaussian distributions, such as Rayleigh, Rician or Nakagami distributions^15,16^. While Gaussian assumptions typically serve as good approximations for GLM-based analysis^17^ and yield high-quality BURST images, more accurate statistical estimation could be achieved by fitting an appropriate distribution to post-collapse frames and using the fitted model for outlier detection on a collapse frame (see **Supplementary Material**).

Accounting for spatiotemporal correlations may further improve estimation of GV signals and associated statistics. Singular value decomposition (SVD)-based filtering^18,19^ can effectively remove coherent background when GVs occupy a small portion of the image, relaxing the constant-background assumption implicit in the template *T* = [1 **0**_1×*N*_] and the estimator 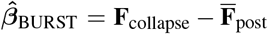 Markov random field models^20,21^ may be employed to encode spatial and/or temporal dependencies, though they may require a large number of background frames. Temporal filtering strategies, such as high-pass filtering, can suppress linear trends or low-frequency components.

The proposed method remains susceptible to motion in the image sequence. While less problematic for relatively static tissues (e.g., the brain of head-fixed mice), it poses challenges for tissues subject to movement (e.g., subcutaneous tumors or the gut). In such cases, image sequences should ideally be acquired during periods of minimal motion – assuming a breathing rate of ∼2 Hz and a frame rate of ∼40 Hz, ∼10 near-static frames may be obtained. If collapse frames are contaminated by motion, or if too few static frames are available, motion correction via image registration or patch-based processing^22^ may be necessary. Alternatively, advances in RF-domain processing^23^ might reduce the number of frames needed for reliable BURST image reconstruction, thereby mitigating motion artifacts.

Finally, when GV collapse spans multiple frames, our framework currently requires manual identification of when collapse signals have fully disappeared. Automatic change-detection algorithms could streamline this step. Such approaches may be especially beneficial in situations when GVs at different depths collapse at different times (e.g., due to shielding by superficial GVs), loosening the overconservative statistical thresholds imposed by multiple-comparison corrections. Taken together, these potential extensions would strengthen the statistical underpinning of BURST analysis and facilitate more robust interpretation of BURST images.

## SUPPLEMENTARY MATERIALS

See the supplementary material for details of the *in vitro* BURST imaging experiment, the distribution of *r*_BURST_ in real image datasets, and statistical BURST processing based on more complex fluctuation models.

## ACKNOWLEDGMENTS

We thank Robin Y. Wen for reviewing our manuscript and its mathematical content, and Shiyu Zhang for helpful discussions. This study was supported by the National Institutes of Health (grants R01EB018975 and R01NS120828 to M.G.S.) and the Chan-Zuckerberg Initiative. M.G.S. is a Howard Hughes Medical Institute Investigator.

## AUTHOR DECLARATION

## Conflict of Interest

M.G.S. is a co-founder of Merge Labs.

## Author Contributions

**Sunho Lee**:Conceptualization (lead); Formal Analysis (lead); Investigation (lead); Methodology (lead); Visualization (lead); Writing – original draft (lead); Writing – review and editing (equal). **Shirin Shivaei**: Conceptualization (supporting); Writing – original draft (supporting); Writing – review and editing (equal). **Mikhail G. Shapiro**: Funding acquisition (lead); Supervision (lead); Writing – original draft (supporting); Writing – review and editing (equal).

## DATA AVAILABILITY

The data that support the findings of this study are available on request from the corresponding author. The code for implementing statistical BURST processing is publicly available on GitHub at https://github.com/shapiro-lab/statistical-BURST.

## Appendix A: Least-squares solution for template-based unmixing matrix equation

### 1. Single collapse frame

Let *M* denote the number of pre-collapse frames and *N* the number of post-collapse frames. The design matrix (**Eq. 1**) is

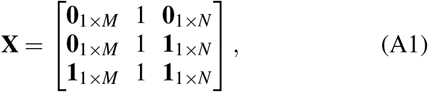

where **0**_*a*×*b*_ and **1**_*a*×*b*_ denote the zero and all-ones matrices of size *a* × *b*, respectively.

Assuming voxel fluctuations are normally distributed with zero means, the maximum-likelihood estimator of **Eq. 2** is equivalent to the ordinary least-squares solution. The closed-form solution is

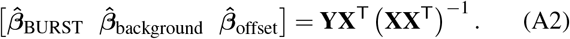

The term **X**^T^(**XX**^T^)^1^ expands to

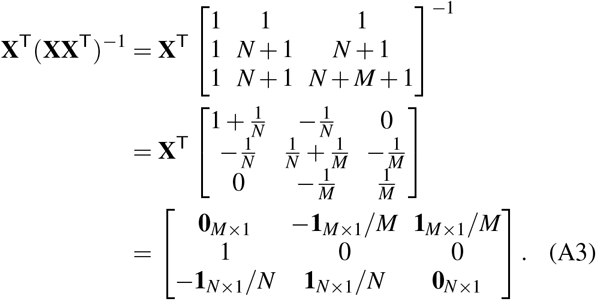

The estimator 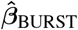is then

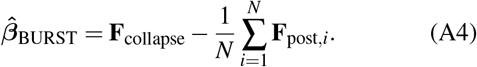

#### 2. Multiple collapse frames

Here, we derive the subtraction form of template-based unmixing for a generalized BURST image sequence in which GV collapse occurs across multiple frames. Suppose collapse is observed for *d* frames upon increasing pressure and treat the signal in each collapse frame as arising from independent GVs. The design matrix then expands to

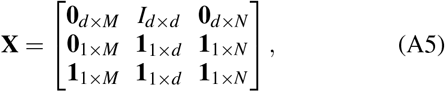

where *I*_*d*_×_*d*_ is the identity matrix of size *d*.

As in **Appendix A 1**, we obtain the closed-form solution by calculating **X**^T^(**XX**^T^)^−1^:

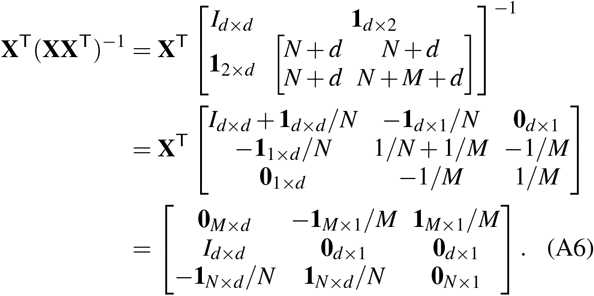

The estimator 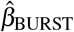for each collapse frame is then

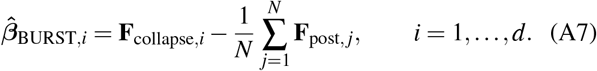

## Appendix B: Equivalence of *r*_BURST_ and *t*_BURST_

Although the equivalence between *r*_BURST_ and *t*_BURST_ can be established directly using the well-known identity for simple linear regression

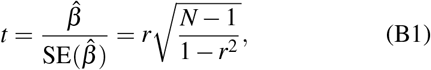

we present here a numerical proof. The mean of all frames 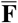can be written as

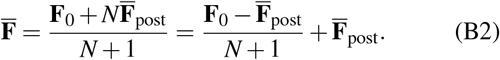

Then, the total variance term in the denominator of *r*_BURSTb_ (**Eq. 7**) is

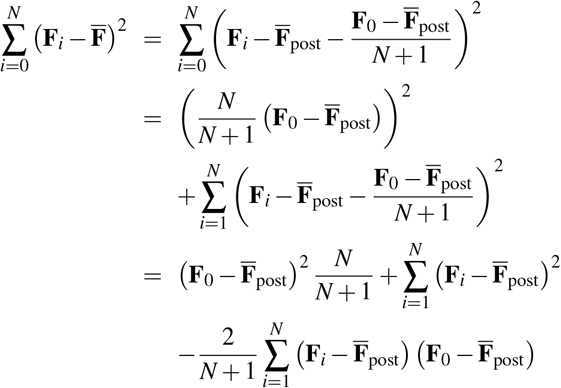

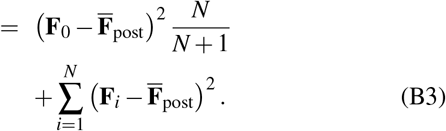

Plugging this into *r*_BURST_ (**Eq. 7**), the correlation becomes

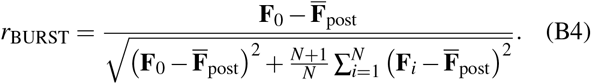

Taking the reciprocal of 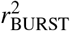

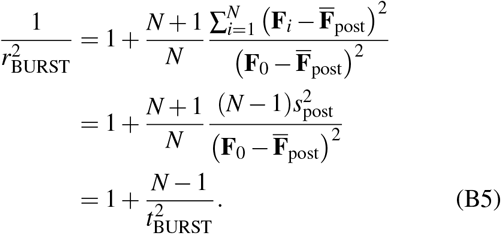

Therefore,

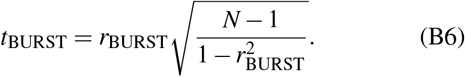

## References

1 A. E. Walsby, “Gas vesicles,” Microbiological Reviews 58, 94–144 (1994).

2 H. Yusefi and B. Helfield, “Ultrasound Contrast Imaging: Fundamentals and Emerging Technology,” Frontiers in Physics 10 (2022).

3 M. G. Shapiro, P. W. Goodwill, A. Neogy, M. Yin, F. S. Foster, D. V. Schaffer, and S. M. Conolly, en”Biogenic gas nanostructures as ultrasonic molecular reporters,” Nature Nanotechnology 9, 311–316 (2014), number: 4.

4 D. Maresca, A. Lakshmanan, A. Lee-Gosselin, J. M. Melis, Y.-L. Ni, R. W. Bourdeau, D. M. Kochmann, and M. G. Shapiro, “Nonlinear ultrasound imaging of nanoscale acoustic biomolecules,” Applied Physics Letters 110, 073704 (2017).

5 D. Maresca, D. P. Sawyer, G. Renaud, A. Lee-Gosselin, and M. G. Shapiro, en”Nonlinear X-Wave Ultrasound Imaging of Acoustic Biomolecules,” Physical Review X 8, 041002 (2018).

6 D. P. Sawyer, A. Bar-Zion, A. Farhadi, S. Shivaei, B. Ling, A. Lee-Gosselin, and M. G. Shapiro, “Ultrasensitive Ultrasound Imaging of Gene Expression with Signal Unmixing,” Nature methods 18, 945–952 (2021).

7 S. Shivaei, A. Liu, M. H. Abedi, J. Revilla, I. U. Hurvitz, M. B. Swift, and M. G. Shapiro, en”Non-invasive imaging of cell-based therapies using acoustic reporter genes,” (2024), pages: 2024.11.01.621111 Section: New Results.

8 S. Shivaei, K. Y. M. Cheung, A. Yadav, I. U. Hurvitz, S. Lee, J. Revilla, C. Rabut, E. Criado-Hidalgo, R. J. Zhang, and M. G. Shapiro, en”Ultrasound imaging of in situ transcriptional activity in opaque tissue,” (2025), iSSN: 2692-8205 Pages: 2025.07.06.663365 Section: New Results.

9 M. T. Buss, L. Zhu, J. H. Kwon, J. J. Tabor, and M. G. Shapiro, en”Probiotic acoustic biosensors for noninvasive imaging of gut inflammation,” Nature Communications 16, 7931 (2025).

10 B. Ling, B. Gungoren, Y. Yao, P. Dutka, R. Vassallo, R. Nayak,C. A. B. Smith, J. Lee, M. B. Swift, and M. G. Shapiro, en”Truly Tiny Acoustic Biomolecules for Ultrasound Imaging and Therapy,” Advanced Materials 36, 2307106 (2024), _eprint: https://onlinelibrary.wiley.com/doi/pdf/10.1002/adma.202307106.

11 R. C. Hurt, M. T. Buss, M. Duan, K. Wong, M. Y. You, D. P. Sawyer,M. B. Swift, P. Dutka, P. Barturen-Larrea, D. R. Mittelstein, Z. Jin, M. H. Abedi, A. Farhadi, R. Deshpande, and M. G. Shapiro, en”Genomically mined acoustic reporter genes for real-time in vivo monitoring of tumors and tumor-homing bacteria,” Nature Biotechnology 41, 919–931 (2023).

12 John F Kenney, engMathematics Of Statistics Part Two (1939).

13 H. Hotelling, “New Light on the Correlation Coefficient and its Transforms,” Journal of the Royal Statistical Society. Series B (Methodological) 15, 193–232 (1953).

14 J. Ahlers, D. Althviz Moré, O. Amsalem, A. Anderson, G. Bokota,P. Boone, J. Bragantini, G. Buckley, A. Burt, M. Bussonnier, A. Can Solak,C. Caporal, D. Doncila Pop, K. Evans, J. Freeman, L. Gaifas, C. Gohlke,K. Gunalan, H. Har-Gil, M. Harfouche, K. I. S. Harrington, V. Hilsenstein, K. Hutchings, T. Lambert, J. Lauer, G. Lichtner, Z. Liu, L. Liu,A. Lowe, L. Marconato, S. Martin, A. McGovern, L. Migas, N. Miller,H. Muñoz, J.-H. Müller, C. Nauroth-Kreß, J. Nunez-Iglesias, C. Pape,K. Pevey, G. Peña-Castellanos, A. Pierré, J. Rodríguez-Guerra, D. Ross,L. Royer, C. T. Russell, G. Selzer, P. Smith, P. Sobolewski, K. Sofiiuk,N. Sofroniew, D. Stansby, A. Sweet, W.-M. Vierdag, P. Wadhwa, M. Weber Mendonça, J. Windhager, P. Winston, and K. Yamauchi, “napari: a multi-dimensional image viewer for Python,” (2023).

15 T. L. Szabo, enDiagnostic Ultrasound Imaging: Inside Out (Academic Press, 2013) google-Books-ID: wTYTAAAAQBAJ.

16 F. Destrempes and G. Cloutier, en”A Critical Review and Uniformized Representation of Statistical Distributions Modeling the Ultrasound Echo Envelope,” Ultrasound in Medicine & Biology 36, 1037–1051 (2010).

17 A. M. Wink and J. B. T. M. Roerdink, eng”BOLD Noise Assumptions in fMRI,” International Journal of Biomedical Imaging 2006, 12014 (2006).

18 S. Lee, D. Wu, D. Malounda, C. Rabut, and M. G. Shapiro, en”Real-time volumetric imaging of cells and molecules in deep tissues with Takoyaki ultrasound,” (2024), pages: 2024.11.14.623368 Section: New Results.

19 A. Bhattacharjee, S. Turner, L. Diao, S. Zhang, and S. Yoon, en”Signal Detection of Point Targets Using Eigen-Images for Super-Resolution Ultrasound Imaging and Gas Vesicle Localization,” (2025), iSSN: 2692-8205 Pages: 2025.07.02.662077 Section: New Results.

20 Q. Liu, M. Sun, C.-C. Li, and R. J. Sclabassi, en”Change Detection in Image Sequence Based on Markov Random Field and Mean Field Theory,”; in enApplied Research in Uncertainty Modeling and Analysis, edited by N. O. Attoh-Okine and B. M. Ayyub (Springer US, Boston, MA, 2005) pp. 115–137.

21 L. Bruzzone and D. Prieto, “Automatic analysis of the difference image for unsupervised change detection ”, IEEE Transactions on Geoscience and Remote Sensing 38, 1171–1182 (2000).

22 J. Boulanger, A. Gidon, C. Kervran, and J. Salamero, en”A Patch-Based Method for Repetitive and Transient Event Detection in Fluorescence Imaging,” PLOS ONE 5, e13190 (2010).

23 P. Frinking, E. Cespedes, J. Kirkhorn, H. Torp, and N. de Jong, “A new ultrasound contrast imaging approach based on the combination of multiple imaging pulses and a separate release burst ”, IEEE Transactions on Ultrasonics, Ferroelectrics, and Frequency Control 48, 643–651 (2001).

