## Supplementary Materials for "Statistical BURST imaging for high-fidelity biomolecular ultrasound"

#### *In vitro* BURST imaging

GVs were prepared following the approach of Lakshmanan *et al.* [1]. Phantoms were prepared in which cylindrical wells contained GVs embedded in 0.5% agarose, while the background matrix was composed of 0.2% 3 $\mu$m $Al_2O_3$ + 1% agarose. For the GV mixture, GV solutions with an initial optical density (OD) of 0.8 at 500 nm were combined with 1% agarose in a 1:1 ratio, yielding a final OD of 0.4. After solidification, thin strips of aluminum foil were manually embedded adjacent to the GV wells. The phantoms were then submerged in a water bath and imaged using a linear probe (Verasonics, L22-14vX) with the BURST imaging sequence [2]. The focal length was set at 7 mm, corresponding to the centers of the GV wells, and the transmit aperture size was 40. The transmit frequency, waveform duty cycle, and number of half-cycles were 15.625 MHz, 0.67, and 3, respectively. Each frame was acquired in AM mode, and the voltages were increased from 1.6 to 20 V. Two pre-collapse frames were recorded, followed by high-pressure acquisitions: 100 frames for **Fig. 2b** and 20 frames for **Fig. 2c**.

#### Note 1: Distribution of $r_{\text{BURST}}$ in real image datasets

Distributions of $r_{\text{BURST}}$ in real datasets can be influenced by multiple factors, including B-mode/AM variations, motion artifacts (even subtle probe micro-movements), voxel-to-voxel dependence, and others. The null distribution of $r_{\text{BURST}}$ for each image dataset can be estimated using only post-collapse frames, excluding the collapse frame (**Fig. S1a-d**). These estimated distributions show that, for some datasets, the theoretical distribution still provides a good fit (**Fig. S1a**), while for others, slight to notable deviations are observed (**Fig. S1b-c**). Such deviations imply that our statistical thresholding may be stricter or looser than the intended $p$-value.

To obtain a more accurate threshold, one approach is to compute the percentile corresponding to the desired $p$-value directly from the background distribution derived from post-collapse frames. Note, however, that the estimated background distribution can vary across post-collapse time points (**Fig. S1d**). Moreover, percentiles at very small $p$-values (e.g., $\sim 1/\#$ of pixels) are determined by only a few samples near the maximum $r_{\text{BURST}}$; estimating thresholds from multiple subsets of post-collapse frames and aggregating the results may therefore improve reliability. Nevertheless, we have observed that this refinement does not drastically alter the quality of the resulting BURST images compared to using the theoretical threshold (**Fig. S1e**). Therefore, while a dataset-specific empirical calibration is possible, the use of theoretical thresholds would remain an effective and computationally efficient default in most cases.

#### Note 2: Statistical BURST processing based on more complex fluctuation models

Considering that B-mode and AM images are computed as the magnitude of complex signals (i.e., ultrasound signal envelope derived from in-phase and quadrature (IQ) data), the voxel amplitude can be modeled using distributions associated with the magnitude of complex random variables and signal envelopes [3,4], such as the Rayleigh, Rician, and Nakagami distributions. Briefly, the Rayleigh distribution arises from the magnitude of a zero-mean circularly symmetric complex Gaussian random variable. The Rician distribution is the non-central generalization of the Rayleigh distribution. The Nakagami distribution is flexible and can represent a variety of statistical behaviors through its shape parameter $m$, including Rayleigh ($m = 1$) and approximations to Rician ($m > 1$). In the following, we construct statistical metrics based on Rayleigh and Nakagami random variables. These models are particularly convenient because their squared values follow Gamma distributions, which simplifies the statistical derivation. In addition, we present the Mahalanobis distance in the complex IQ plane as an alternative statistical metric.

##### 1. Nakagami case

Assume that, for a background voxel $j$, the envelope amplitudes $F^j_{0,\dots,N}$ are independent and identically distributed as Nakagami$(m,\Omega)$, where $m$ and $\Omega$ denote the shape and scale parameters, respectively. The squared amplitudes $P^j = (F^j)^2$ then follow a Gamma distribution

$$P^j \mid j \in \text{background} \sim \text{Gamma}(m,\theta), \qquad \theta = \frac{\Omega}{m}.$$

We define the test statistic $R_{\text{BURST}}$ (**Fig. S2a**) as the ratio of the squared collapse frame (i.e., the power at the collapse frame) to the average of the squared post-collapse frames (i.e., the average power across post-collapse frames):

$$R^j_{\text{BURST}} \equiv \frac{P^j_0}{\frac{1}{N}\sum_{i=1}^{N} P^j_i} = \frac{P^j_0}{\bar{P}^j_{\text{post}}}.$$

Because $\sum_{i=1}^{N} P^j_i \sim \text{Gamma}(Nm, \theta)$ and is independent of $P^j_0$, $R^j_{\text{BURST}}$ follows an $F$-distribution with $2m$ and $2Nm$ degrees of freedom:

$$R^j_{\text{BURST}} \mid j \in \text{background} \sim F_{2m,2Nm}.$$

Since the distribution depends on the unknown shape parameter $m$, it must be estimated from the post-collapse frames. The maximum-likelihood estimate $\hat{m}$ satisfies[5]

$$\ln(\hat{m}) - \psi(\hat{m}) = \ln(\overline{P}_{\text{post}}) - \overline{\ln P_{\text{post}}},$$

where $\psi$ is the digamma function and $\overline{\ln P_{\text{post}}} = \frac{1}{N}\sum_{i=1}^{N} \ln P_i$. This equation can be solved via numerical approaches such as the Newton–Raphson method[6]. A faster alternative is the method-of-moments estimate[5]

$$\hat{m}_{\text{mom}} = \frac{\overline{P}^2_{\text{post}}}{\text{Var}(P_{\text{post}})},$$

where $\text{Var}(P_{\text{post}})$ is the sample variance of the squared post-collapse frames. The estimation of $m$—and thus the performance of the Nakagami-based ratio test—generally improves as the number of post-collapse frames increases.

#### 2. Rayleigh case

Because the Rayleigh distribution arises from a zero-mean complex random variable, it can be beneficial to remove a persistent (non-zero-mean) background component in the complex IQ domain before taking magnitude images. Analogous to **Eq. 3**, we subtract the average of the post-collapse IQ frames (i.e., estimated background) from all IQ frames,

$$\mathbf{F}_{(\cdot)} = \left|\mathbf{IQ}_{(\cdot)} - \frac{1}{N}\sum_{i=1}^{N}\mathbf{IQ}_i\right| = \left|\mathbf{IQ}_{(\cdot)} - \overline{\mathbf{IQ}}_{\text{post}}\right|.$$

Alternatively, singular value decomposition (SVD)-based filtering[7] can be used to suppress the temporally consistent background component.

As in the Nakagami case, we can work with squared images since the square of a Rayleigh random variable also follows a Gamma distribution. Because Rayleigh is a special case of Nakagami with $m = 1$, we use the same statistic $R_{\text{BURST}}$, which under the background hypothesis follows

$$R^j_{\text{BURST}} \mid j \in \text{background} \sim F_{2,2N}.$$

Notably, in the Rayleigh case, the null distribution of $R_{\text{BURST}}$ depends only on the number of post-collapse frames $N$, similar to $t_{\text{BURST}}$ (**Eq. 11**). Users may still apply the Nakagami ratio test after IQ processing since Nakagami includes the Rayleigh case, but this reintroduces the need to estimate $m$ for each pixel.

A one-sided $p$-value for testing $P_0 > \overline{P}_{\text{post}}$ under $F_{2,2N}$ simplifies to a closed-form expression directly computable from the squared frames:

$$p = 1 - I_{\frac{R_{\text{BURST}}}{R_{\text{BURST}}+N}}(1,N) = \left(\frac{\sum_{i=1}^{N} P_i}{\sum_{i=0}^{N} P_i}\right)^N,$$

where $I$ is the regularized incomplete beta function[8,9].

#### 3. Mahalanobis distance

In the Rayleigh case, after subtracting the temporally consistent background component, we assumed that the residual complex IQ signal at a background pixel was circularly symmetric. A more general model is to assume that, at each background pixel $j$, the real and imaginary parts of the IQ signal are jointly Gaussian. Writing the complex IQ value as the 2D real vector

$$\mathbf{z}^j_{(\cdot)} = \begin{bmatrix} \Re\left(IQ^j_{(\cdot)}\right) \\ \Im\left(IQ^j_{(\cdot)}\right) \end{bmatrix}$$

and assuming

$$\mathbf{z}^{j}_{(\cdot)} \mid j \in \text{background} \sim \mathcal{N}_2\left(\boldsymbol{\mu}^{j}, \Sigma^{j}\right),$$

where $\boldsymbol{\mu}^{j}$ and $\Sigma^{j}$ are the mean vector and covariance matrix, respectively. We then quantify how unusual the collapse-frame IQ value is relative to the post-collapse distribution using the (squared) Mahalanobis distance[10] (**Fig. S2a**):

$$d_M^2 = \left(\mathbf{z}_0^j - \bar{\mathbf{z}}_{\text{post}}^j\right)^\top \left(S_{\text{post}}^j\right)^{-1} \left(\mathbf{z}_0^j - \bar{\mathbf{z}}_{\text{post}}^j\right),$$

where $\bar{\mathbf{z}}_{\text{post}}^j$ and $S_{\text{post}}^j$ are the sample mean and sample covariance of the post-collapse IQ values, respectively. With appropriate scaling, this becomes Hotelling's two-sample $T^2$ statistic (with sample sizes 1 and $N$)[11]:

$$T^2 = \frac{N}{N+1} d_M^2.$$

Under the background hypothesis, the $T^2$ statistic can be related to the $F$-distribution by

$$\frac{N-2}{2(N-1)} T^2 \sim F_{2,N-2}.$$

Therefore, the null distribution of the Mahalanobis distance is

$$d_M \sim \sqrt{\frac{2(N-1)(N+1)}{N(N-2)} F_{2,N-2}}.$$

The image results obtained using all statistical tests proposed above are shown in **Fig. S2b**. In practice, we generally observed only subtle differences among the various approaches (including the Gaussian case)—much smaller than the improvement gained by using a statistical method versus the conventional method. In addition, IQ processing occasionally made some GV pixels more apparent that were not clearly visible beforehand. We do not attempt to determine which statistical test is optimal, nor to fully characterize the effects of IQ processing, as these topics are beyond the scope of this paper. Instead, we include one representative *in vitro* example and provide implementations of all methods in our MATLAB script so users can choose the approach most suitable for their applications.

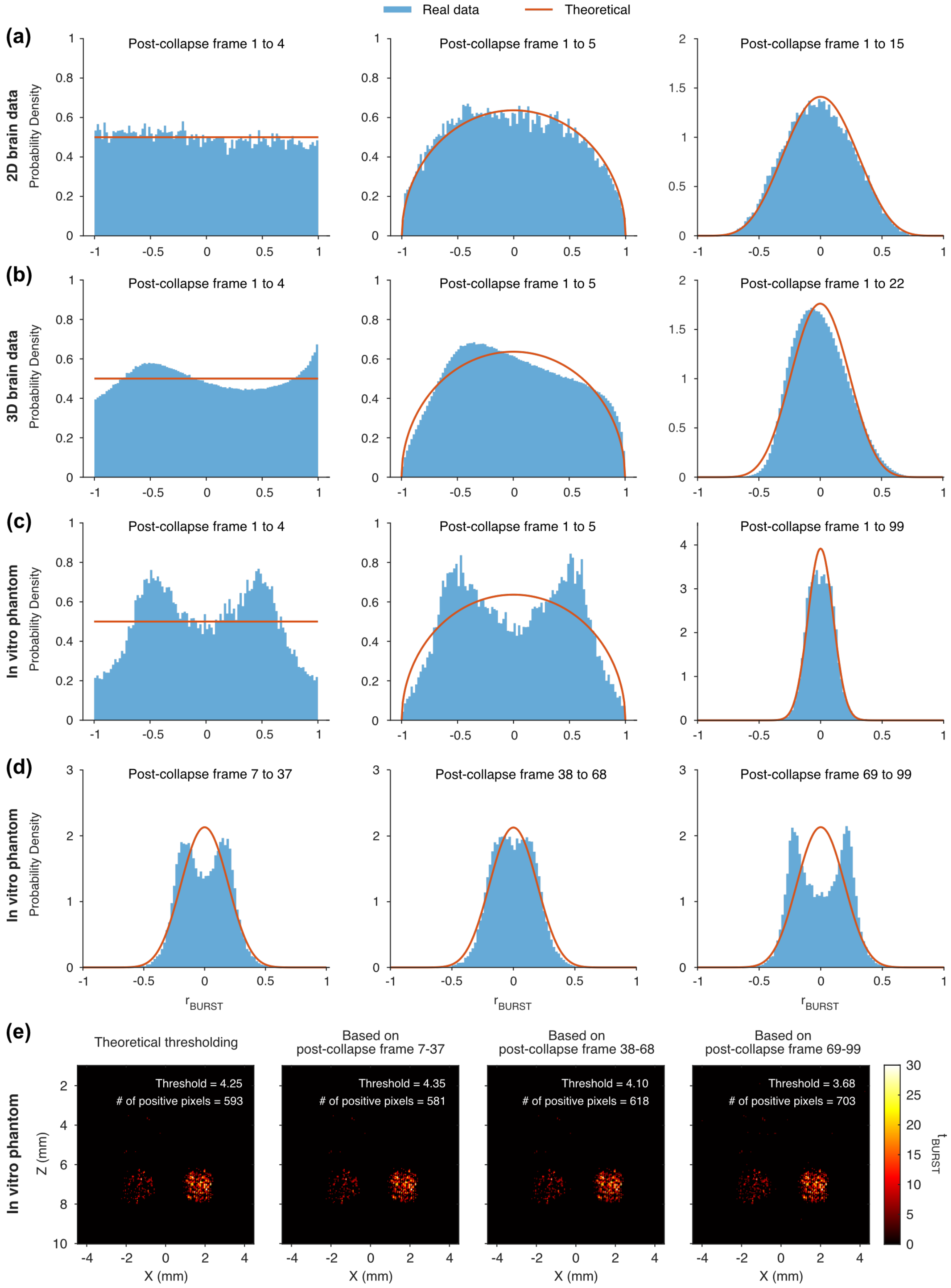


Figure S1: **Background distribution of $r_{BURST}$ from real datasets.** The background $r_{BURST}$ values are calculated using only post-collapse frames. Blue bars and red lines represent the histogram of $r_{BURST}$ and its theoretical PDF, respectively. (a) 2D brain data from **Fig. 1**, (b) 3D brain data from *Shivaei et al.*[12] and (c-e) *in vitro* phantom data from **Fig. 2b** are used. (e) The $t_{BURST}$ map of the *in vitro* phantom image is thresholded using either a theoretical value or a percentile of the background distribution obtained from post-collapse frames corresponding to $p = 1 \times 10^{-4}$. The threshold values and the number of positively classified pixels are shown at the top right of each image. The number of pixels is (X, Z) = 178 $\times$ 183.

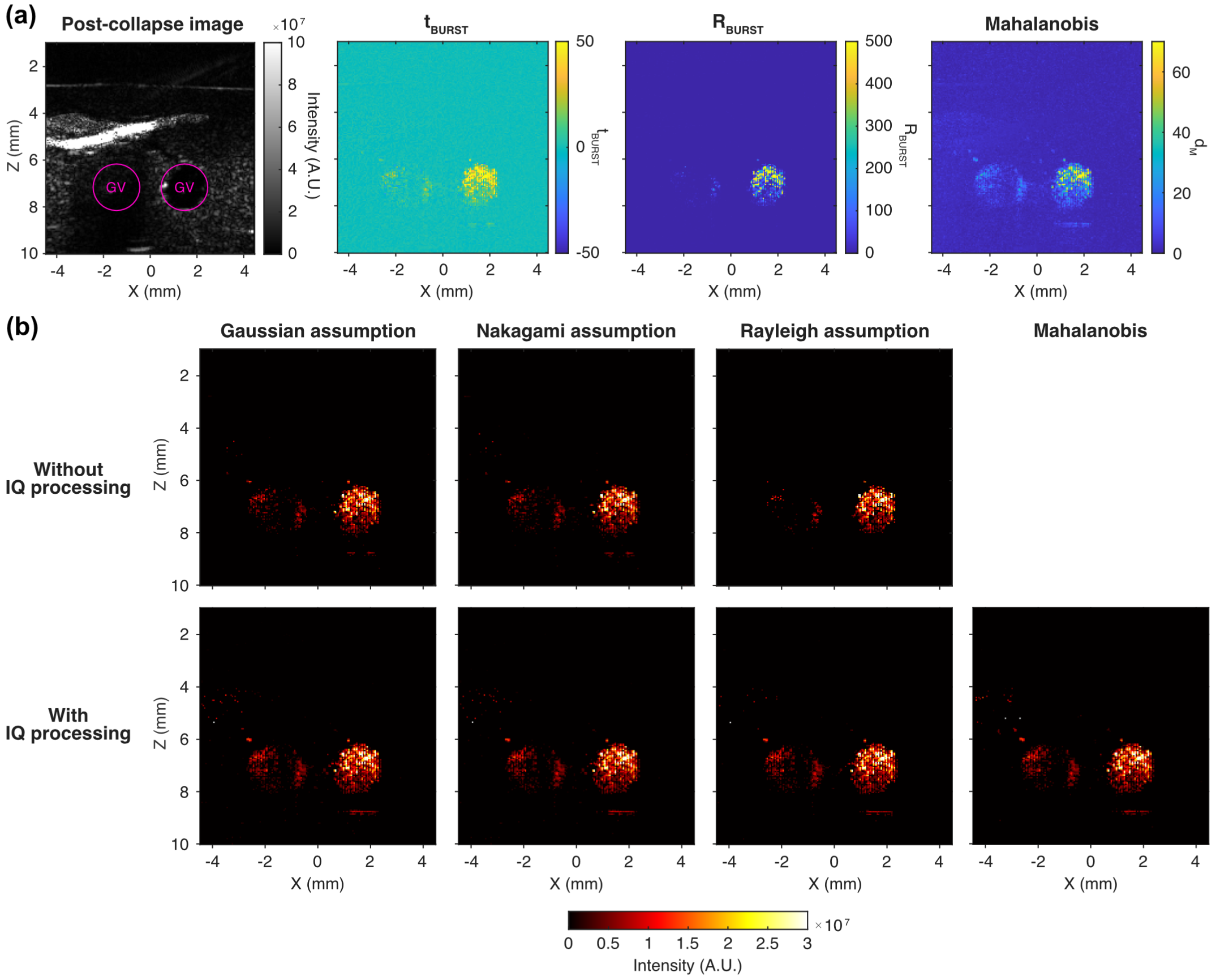


Figure S2: **Statistical BURST processing under different fluctuation assumptions.** (a) Statistical parameter maps ($t_{\mathrm{BURST}}$, $R_{\mathrm{BURST}}$, and Mahalanobis distance) for the *in vitro* example in **Fig. 2c**. The last post-collapse frame is included for structural reference. IQ processing was not performed when generating $t_{\mathrm{BURST}}$ and $R_{\mathrm{BURST}}$. (b) The BURST image generated without IQ processing under a Gaussian assumption (top left) corresponds to the primary method used in this study. The phantom is the same as **Fig. 2c**, but IQ frames were used here. IQ processing was performed by subtracting the average post-collapse IQ frame from each IQ frame and/or by calculating the Mahalanobis distance. BURST images generated using thresholded statistical maps based on Nakagami-based (second column), Rayleigh-based (third column), and Mahalanobis-distance-based (rightmost column) statistics are also shown. All images used $p = 1 \times 10^{-4}$. For the intensity map, $\beta_{\mathrm{BURST}} = \mathbf{F}_0 - \overline{\mathbf{F}}_{\mathrm{post}}$ was computed either without IQ processing (top row; $\mathbf{F}_{(\cdot)} = |\mathbf{IQ}_{(\cdot)}|$) or with IQ processing (bottom row; $\mathbf{F}_{(\cdot)} = |\mathbf{IQ}_{(\cdot)} - \overline{\mathbf{IQ}}_{\mathrm{post}}|$).

### References


[1] Anupama Lakshmanan, George J. Lu, Arash Farhadi, Suchita P. Nety, Martin Kunth, Audrey Lee-Gosselin, David Maresca, Raymond W. Bourdeau, Melissa Yin, Judy Yan, Christopher Witte, Dina Malounda, F. Stuart Foster, Leif Schröder, and Mikhail G. Shapiro. Preparation of biogenic gas vesicle nanostructures for use as contrast agents for ultrasound and MRI. *Nature Protocols*, 12(10):2050–2080, October 2017.

[2] Daniel P. Sawyer, Avinoam Bar-Zion, Arash Farhadi, Shirin Shivaei, Bill Ling, Audrey Lee-Gosselin, and Mikhail G. Shapiro. Ultrasensitive Ultrasound Imaging of Gene Expression with Signal Unmixing. *Nature methods*, 18(8):945–952, August 2021.

[3] François Destrempes and Guy Cloutier. A Critical Review and Uniformized Representation of Statistical Distributions Modeling the Ultrasound Echo Envelope. *Ultrasound in Medicine & Biology*, 36(7):1037–1051, July 2010.

[4] Michael L. Oelze and Jonathan Mamou. Review of quantitative ultrasound: envelope statistics and backscatter coefficient imaging and contributions to diagnostic ultrasound. *IEEE transactions on ultrasonics, ferroelectrics, and frequency control*, 63(2):336–351, February 2016.

[5] Athanasios Papoulis and S. Unnikrishna Pillai. *Probability, Random Variables, and Stochastic Processes*. McGraw-Hill, 4th edition, 2002. Google-Books-ID: cUmiDAEACAAJ.

[6] S. C. Choi and R. Wette. Maximum Likelihood Estimation of the Parameters of the Gamma Distribution and Their Bias. *Technometrics*, 11(4):683–690, November 1969. _eprint: https://www.tandfonline.com/doi/pdf/10.1080/00401706.1969.10490731.

[7] Charlie Demené, Thomas Deffieux, Mathieu Pernot, Bruno-Félix Osmanski, Valérie Biran, Jean-Luc Gennisson, Lim-Anna Sieu, Antoine Bergel, Stéphanie Franqui, Jean-Michel Correas, Ivan Cohen, Olivier Baud, and Mickael Tanter. Spatiotemporal Clutter Filtering of Ultrafast Ultrasound Data Highly Increases Doppler and fUltrasound Sensitivity. *IEEE transactions on medical imaging*, 34(11):2271–2285, November 2015.

[8] Stephen Wolfram. *The Mathematica Book*. Wolfram Media Inc, Champaign, Ill., 5th edition, 2003.

[9] DLMF: §8.17 Incomplete Beta Functions Related Functions Chapter 8 Incomplete Gamma and Related Functions.

[10] Reprint of: Mahalanobis, P.C. (1936) "On the Generalised Distance in Statistics.". *Sankhya A*, 80(1):1–7, December 2018.

[11] Kantilal V. Mardia, John T. Kent, and John M. Bibby. *Multivariate analysis*. Probability and mathematical statistics a series of monographs and textbooks. Acad. Press, Amsterdam, tranferred to digital pr edition, 2006.

[12] Shirin Shivaei, Kathy Y. M. Cheung, Akanksha Yadav, Isabella U. Hurvitz, Sunho Lee, Julio Revilla, Claire Rabut, Ernesto Criado-Hidalgo, Raymond J. Zhang, and Mikhail G. Shapiro. Ultrasound imaging of in situ transcriptional activity in opaque tissue, July 2025. ISSN: 2692-8205 Pages: 2025.07.06.663365 Section: New Results.
